# Alpha oscillations are dysrhythmic in Fragile X syndrome

**DOI:** 10.64898/2026.01.30.702828

**Authors:** Peyton Siekierski, Yanchen Liu, Grace Westerkamp, Rana Elmaghraby, Zag ElSayed, Donald Gilbert, Craig A. Erickson, Ernest V. Pedapati

## Abstract

**Background:** Alpha oscillations are dominant rhythms in the human brain, supporting inhibitory control and coordination of neural activity. Altered alpha dynamics are observed across many neuropsychiatric and neurodevelopmental disorders, including Fragile X syndrome (FXS), the most common monogenic cause of autism and intellectual disability. FXS exhibits paradoxical alpha power: elevated absolute but reduced relative power. To resolve this incongruity, we considered that conventional power metrics, relying on averaging, may obscure the underlying critical temporal dynamics of such neural rhythms.

**Methods:** Here, we investigate alpha oscillations in FXS as a model to decompose nonspecific alpha abnormalities into underlying temporal features. We used cycle-by-cycle (*bycycle*) alpha burst analysis from source-localized resting-state EEG of 70 individuals with FXS (20.5±10.0 years; 32 females, 38 males) and 71 age- and sex-matched typically developing controls (22.2±10.7 years; 30 females, 41 males). Statistical modeling examined group, sex, and regional differences in alpha burst features using generalized linear mixed-effects models.

**Results:** We reveal that alpha bursts in FXS show reduced count only in males, prolonged periods across sexes, and elevated amplitudes, particularly in males. Spatial mapping identified differential circuit vulnerability: timing-associated dysregulation in cognitive-control regions and amplitude elevations in sensory cortices. Within the FXS group, global alpha burst amplitude correlated with hyperactivity symptoms and inversely with general intelligence scores, and burst count correlated with age.

**Limitations:** This study is limited by its resting-state design and cross-sectional nature. Future studies should explore task-based modulation of alpha burst features and longitudinal trajectories in FXS. Additionally, fragile X messenger ribonucleoprotein (FMRP) was not quantified for participants, limiting potential stratification by molecular severity.

**Conclusions:** These findings resolve paradoxical alpha power in FXS into features consistent with interneuron dysfunction, demonstrating the potential for burst-level decomposition in mechanistic hypothesis generation and biomarker development across neurodevelopmental and neuropsychiatric disorders.

## Introduction

Alpha oscillations (8–12.5 Hz) are a prominent and functionally significant oscillation in the human brain and have been implicated as a mechanism for inhibitory control and timing of neural activity^1, 2^. Alpha oscillation generation and propagation is complex and incompletely understood, with the prevailing view implicating thalamocortical and cortico-cortical circuits, suggesting that alpha dysfunction may manifest differently across brain regions^3–7^. Alterations in alpha oscillations are common across clinical populations and reflect a shared transdiagnostic vulnerability. Peak alpha frequency (PAF) is diminished in autism spectrum disorder (ASD), psychotic disorders, and Parkinson’s disease^8–10^. Alpha disruptions linked with changes in high-frequency activity (termed thalamocortical dysrhythmia) have been identified in epilepsy, schizophrenia, and major depressive disorder^11–15^. Fragile X syndrome (FXS) exemplifies this motif, exhibiting reduced relative alpha power, slowed alpha frequency, increased gamma power, and abnormal theta–gamma coupling^16–19^. Notably, reduced relative alpha power in FXS correlates with symptom severity, including increased anxiety, decreased auditory attention, and impaired social functioning^16^. FXS offers a genetically defined model to study changes in alpha activity which may reflect compromised network coordination and timing^20^.

FXS provides an ideal model to dissect alpha dysfunction mechanisms, as the most common monogenic cause of intellectual disability and autism. This disorder is caused by the silencing of the *FMR1* gene, leading to loss of fragile X messenger ribonucleoprotein (FMRP). FMRP is a key regulator of dendritic maturation, synaptic protein synthesis, and experience-dependent circuit refinement^21–24^. X-linked inheritance results in high clinical penetrance and typically greater phenotypic severity in males due to near-complete FMRP absence, whereas affected females are obligate mosaics with variable FMRP expression due to random X-inactivation.

Individuals affected by FXS manifest impairments in cognition, language, and social interactions, leading to lifelong disability^25^. Thus far, pharmacological interventions that have shown efficacy in *Fmr1* knockout (KO) mouse models have largely failed to translate to meaningful clinical benefits in humans^26–28^. This gap reflects our limited understanding of how FMRP loss uniquely disrupts neural function in humans and a misalignment with translational biomarkers, which represent a key obstacle to therapeutic progress in FXS^29^. Alpha oscillations represent a prime candidate to explain the discrepancy between humans and rodent models: in human FXS, these rhythms show alterations that correlate with clinical measures^16, 30^, yet they remain poorly characterized in *Fmr1* KO mouse models^31–33^.

Interestingly, power spectrum analysis of the alpha band in FXS has paradoxically found reduced relative alpha power yet largely elevated absolute alpha power^16^ (**Fig. 1**). We can reconcile this contradiction methodologically as conventional Fourier-based approaches time-average neural activity and thus overlook the dynamic, burst-like nature of neural oscillations^34, 35^. Furthermore, these approaches obscure raw waveform features that may provide deeper insight into the underlying substrate which generate coherent rhythmic field potentials from vast neuronal populations^36^. Cycle-by-cycle analysis (“bycycle”) represents a methodological advancement that preserves the time-related structure of neural oscillations while extracting detailed waveform characteristics for individual oscillatory cycles^37^. This approach enables quantification of alpha burst detection, amplitude modulation, and period variability in the time domain, providing a more nuanced window into compound alpha oscillatory dysfunction. Cycle-by-cycle analysis complements other time-resolved approaches such as empirical mode decomposition and burst detection algorithms used to characterize transient oscillations in movement disorders and attention paradigms^38, 39^.

**Fig. 1.**
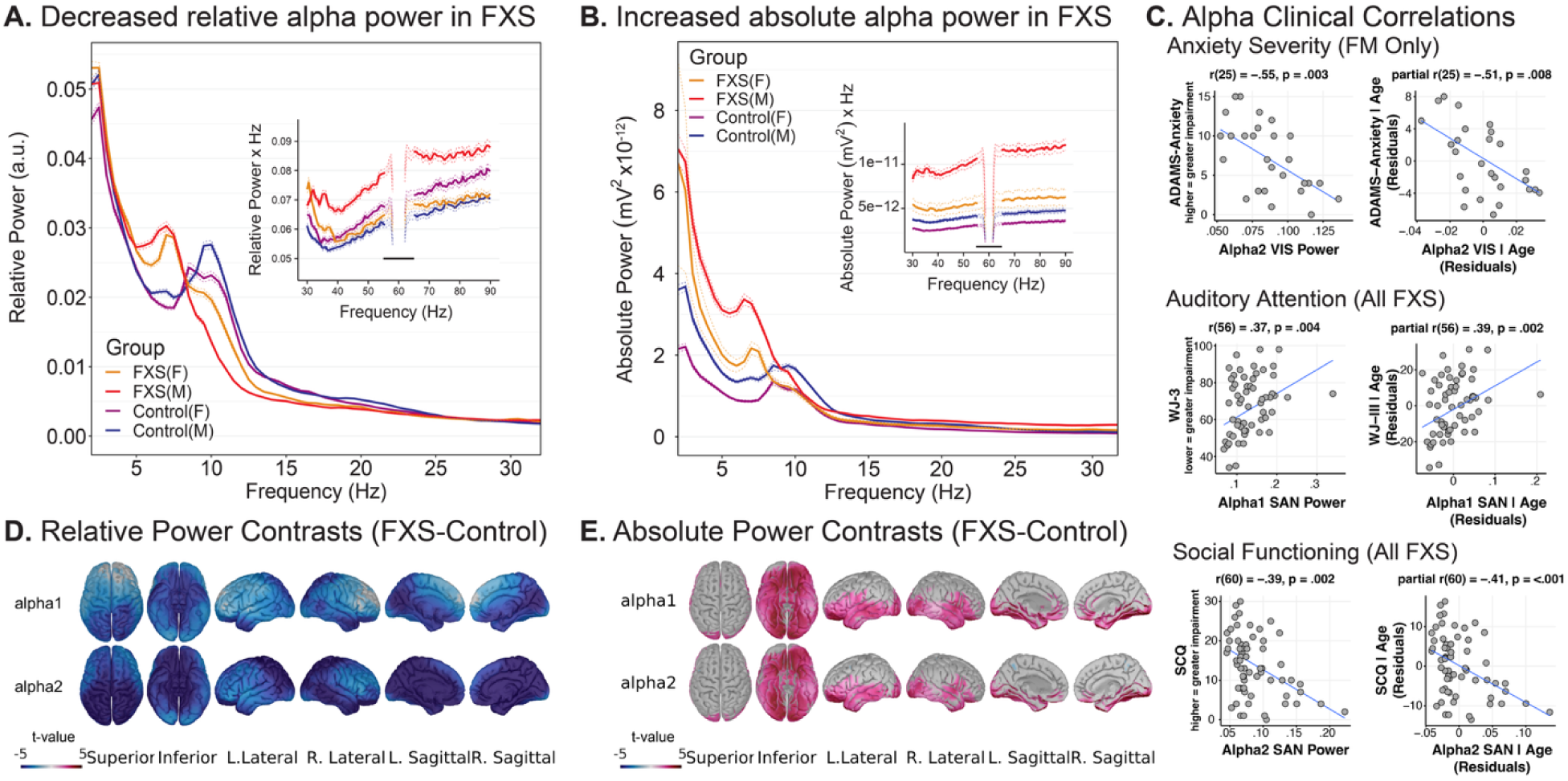
Decreased relative and increased absolute alpha power in FXS. (a–b) Scalp EEG power spectra for males and females with FXS and age- and sex-matched controls demonstrate a paradox between relative and absolute power. **(a)** Group-averaged relative power spectra (solid lines ± 95% CI) show reduced alpha-band power (8-12.5 Hz) and a leftward shift of the dominant peak frequency in FXS. **(b)** In contrast, absolute power spectra reveal elevated alpha and broadband power in FXS groups. Insets highlight high-frequency (30–90 Hz) gamma activity. **(c)** Significant correlations between alpha power and clinical measures across FXS participants and a subgroup of full-mutation (FM) males. Scatterplots show subject-level bivariate and age-corrected partial Spearman correlations, illustrating associations between alpha power and anxiety severity, auditory attention, and social functioning. **(d–e)** Source-level group contrasts (FXS – Control) illustrate the spatial distribution of these effects, with widespread decreases in relative alpha power **(d)** and corresponding increases in absolute alpha power **(e)**. Warmer colors indicate FXS > control, cooler colors FXS < control (non-significant vertices shown in gray). Adapted from Pedapati et al., 2022 (Fig. 2b–d, 5d, Supplementary Fig. 2), which reported the original spectral analyses of this same dataset; panels were reformatted for presentation. Licensed under Creative Commons Attribution 4.0 International License.

This study leverages a well-powered previously published resting-state EEG dataset^16^ to conduct a comprehensive cycle-by-cycle analysis of alpha oscillations in FXS. We conducted our analysis following EEG source localization^40^ to characterize distribution of alpha burst disruptions across space and time. We hypothesized that individuals with FXS would exhibit specific alterations in burst features consistent with their paradoxical power findings: reduced burst occurrence (potentially explaining reduced relative power), prolonged cycle periods (consistent with frequency slowing), and increased burst amplitude (potentially explaining elevated absolute power). Additionally, we anticipated sex-specific patterns given FMRP expression differences between males and females with FXS. These considerations implicate aberrant alpha dynamics as a key feature of FXS pathophysiology and highlight the need for time domain-associated approaches to reveal their underlying mechanisms.

## Methods and Materials

### Sample

Resting-state EEG recordings from a previously published dataset from our laboratory^16^, comprised of individuals with FXS (n=70, 32 females and 38 males; 20.5±10.0 years) and age-and sex-matched TDC (n=71, 30 females and 41 males; 22.2±10.7 years) (National Institutes of Mental Health U54 HD082008). All participants provided written informed consent (or assent, where applicable), and the study was approved by the institutional review board of Cincinnati Children’s Hospital Medical Center. Clinical characteristics of each cohort are presented in Table 1. TDC participants with any neuropsychiatric condition or current medication use were excluded. FXS participants were excluded if there were taking benzodiazepines and/or antiepileptics. FXS participants taking antidepressants, atypical antipsychotics, or other concurrent medications were included, provided they had been on a stable dose for at least six weeks prior to enrollment. This approach reflects typical medication use patterns observed in individuals with FXS.

**Table 1.** Demographics and clinical variables.

| Characteristic | FXS, N = 70<sup>1</sup> | TDC, N = 71<sup>1</sup> | p-value<sup>2</sup> |
| --- | --- | --- | --- |
| Sex |  |  |  |
| F | 32 (46%) | 30 (42%) |  |
| M | 38 (54%) | 41 (58%) |  |
| Age at Visit | 20.5±10.0 | 22.2±10.7 | 0.35 |
| FSIQ | 49.5±30.0 | 103.2±9.2 | <0.001 |
| NVIQ | 40.0±35.7 | 103.4±10.7 | <0.001 |
| VIQ | 58.9±28.7 | 103.1±12.8 | <0.001 |
| SCQ | 13.6±7.7 | 2.1±2.2 | <0.001 |
| WJ-III | 67.8±15.6 | 93.5±12.0 | <0.001 |
| ADAMS-Depressed | 3.6±4.6 | 1.4±3.1 | <0.001 |
| ADAMS-Social | 6.6±5.3 | 1.3±2.6 | <0.001 |
| ADAMS-Anxiety | 7.2±5.0 | 2.1±2.5 | <0.001 |
| ADAMS-OCD | 2.3±2.5 | 0.4±1.1 | <0.001 |
| ABC FXS subscale 4: Hyperactivity/noncompliance | 9.0±7.5 | 0.8±1.7 | <0.001 |
| ABC FXS subscale 5: Inappropriate speech | 4.5±3.5 | 0.1±0.5 | <0.001 |
| <sup>1</sup>n (%); Mean±SD |  |  |  |
| <sup>2</sup>independent Student t tests, p unadjusted significance |  |  |  |

*FXS* Fragile X Syndrome, *F* female, *M* male, *FSIQ* Full Scale IQ, *NVIQ* Non-verbal intelligence scale, *VIQ* verbal intelligence scale, *SCQ* Social Communication Questionnaire, *WJ-III* Woodcock III Tests of Cognitive Abilities, *ADAMS* Anxiety, Depression, and Mood Scale, *ABC* Aberrant Behavior Checklist

### Data acquisition and preprocessing

EEG data were collected at a 1000 Hz sampling rate using a 128-channel HydroCel electrode net and EGI NetAmp 400 (Magstim/EGI, Eugene, OR). Continuous EEG data were collected for 5 minutes during resting-state conditions with participants seated comfortably while watching a silent video to encourage cooperation. All EEG data were blinded for preprocessing and analysis. Data was exported in EGI raw format and imported into EEGLAB SET format in MATLAB (version 2018b, The MathWorks Inc., Natick, MA, USA). Raw recordings were high pass digital zero-phase filtered at 2 Hz and notch filtered at 55-65 Hz to remove line noise using EEGLAB 14.1.2^41^. Filtered data were visually inspected to remove noisy channels and segments of data. Removed channels (less than 5% per subject) were interpolated using spherical spline interpolation. Independent component analysis (ICA) decomposition was performed using the extended INFOMAX algorithm in EEGLAB and myogenic contamination was reduced^42, 43^. The resulting components corresponding to eye, muscle, and channel artifacts were manually identified and removed. Data were average referenced, down sampled to 500 Hz, and divided into 2-second epochs. To ensure consistent data length across participants for comparative analysis, all recordings were normalized to the shortest recording duration identified across the sample (86 seconds). This approach ensured equal data contributions from each participant.

### Source estimation

Cycle-by-cycle analysis was performed on source-localized time series. For each subject, 86 seconds of artifact-free data from each of the EGI 128-channel electrodes were co-registered with Montreal Neurological Institute (MNI) averaged ICBM152 common brain template^44^. Source localization was performed on blinded artifact-free preprocessed data using a weighted minimum norm estimate (MNE) model to generate a current source density map (units: picoampere-meter; pA-m) in Brainstorm^45^. OpenMEEG was used to compute a 15,000 vertices lead-field mesh incorporating electrode distances^46^. Individual vertices from the lead-field mesh were grouped into 68 cortical nodes according to the Desikan-Killiany (DK) atlas^47^ (Supplementary Fig. 1). The anatomical configuration derived from the DK atlas consisted of categorizing the 68 nodes into 14 regions: prefrontal, frontal, central, temporal, parietal, cingulate, and occipital brain regions with bilateral hemisphere (right (R) or left (L)) representation.

### Burst parameter optimization and burst identification

Oscillatory burst detection was performed using the *bycycle* algorithm, which analyzes neural signals cycle by cycle to identify features of sustained rhythmic activity in the time domain. This method can effectively characterize the individual oscillatory cycles in non-averaged, filtered data^37^ (https://github.com/bycycle-tools/bycycle). Analysis focused on the alpha frequency band (8-12.5 Hz). Bursts consist of a sequence of cycles defined by a set of detection parameters.

Excluding the monotonicity parameter, the standard burst detection thresholds were applied to all subjects: (1) amplitude fraction of 0.3, (2) amplitude consistency of 0.4, (3) period consistency of 0.5, (4) minimum cycle requirement of 3^37^. The monotonicity threshold was specifically optimized for alpha separately to allow for detection of more bursts while maintaining shape constraints for genuine oscillatory activity (Supplementary Fig. 2). Monotonicity threshold of 0.55 was used for alpha. Narrowband filtering parameters included 0.5-second filter windows with automatic cycle determination for the alpha frequency range. Burst detection utilized the ’trough’ as the center extrema reference point, with cycle boundaries defined from trough-to-trough intervals. The standard ’cycles’ burst method was employed to identify sustained periods of oscillatory activity meeting the specified amplitude, consistency, and shape criteria^37^.

### *Bycycle* feature generation

For each detected cycle, the *bycycle* algorithm computed comprehensive waveform characteristics including period, amplitude, and burst count. Period refers to the duration of individual cycles (trough-to-trough intervals). Extracted period values were stored in samples and converted to milliseconds (ms) by dividing by the sampling rate (500 Hz), such that each sample represents 2 milliseconds (1 sample = 2 ms). Amplitude is defined by the peak-to-trough amplitude within each cycle. Burst count is calculated by summing the binary indicator (is_burst) that determines whether each cycle met burst criteria defined above across each normalized recording duration of concatenated epochs.

### Statistical modeling

Generalized linear mixed-effects models (GLMMs) were conducted to account for the hierarchical structure of nodes nested within participants. Models were implemented using the glmmTMB package in R, with appropriate distribution families selected based on the nature of dependent variables (Gaussian for continuous features, Poisson for count data). For each feature, models were built using a forward stepwise approach, beginning with a base model containing only group effects and progressively adding complexity. Note that for burst period, statistical models were fitted using period values in samples, and model estimates were converted to milliseconds for reporting. For burst amplitude, values were log10-transformed (base-10 logarithm) for statistical modeling to normalize the distribution. The modeling process started with group as the sole fixed effect, then sequentially evaluated the addition of sex, region, and their interactions (group × sex, group × region, etc.), retaining terms only when they produced statistically significant improvements to model fit. This iterative approach continued until no further significant effects or interactions could be identified, resulting in the final model specifications (Supplementary Table 1). The final models varied in complexity depending on the specific feature analyzed, ranging from simple main effects models to more complex specifications including: (1) fixed effects for group, sex, and brain region with significant two-way and three-way interactions; (2) random intercepts for participants (1|eegid), participant-region interactions (1|eegid:region), and random intercepts for nodes nesting in region (1|region:node). Post hoc contrasts were conducted using estimated marginal means (emmeans) to evaluate main effects and interactions, with Benjamini-Hochberg corrections applied to control the false discovery rate across multiple pairwise comparisons within each effect. Primary hypotheses tested group differences in oscillatory burst properties, with secondary analyses examining sex differences and regional specificity. All analyses were performed using R statistical software.

Statistical significance was set at α = 0.05, with appropriate corrections applied for multiple testing where applicable. Effects sizes were calculated with 95% confidence intervals using Cohen’s d for continuous outcomes (burst amplitude, burst period), and rate ratio (RR) for Poisson-distributed count data (burst count). Complete effect size calculation results reported in Supplementary Tables 4 and 5.

### Clinical associations

Hierarchical linear regression models were used to identify significant relationships with clinical variables and cycle features while controlling for potential confounding variables. In contrast to the LME analyses which contained a large number of within-participant replicates, our regression models required averaging cycle features per participant, reducing data redundancy. Thereby, outliers were identified and removed when exceeding 3 standard deviations from the mean, resulting in the exclusion of one participant for burst amplitude analyses, one for burst count analyses, and two for period analyses. Models included interaction terms to test for sex differences (clinical measure ∼ cycle feature * sex + visit age), with follow-up sex-stratified analyses. Models were fitted using ordinary least squares regression and conducted at whole-brain level. Model performance was evaluated using adjusted R-squared values, and effect sizes were calculated using Cohen’s f². Statistical significance was assessed at p < 0.05. All analyses were conducted in R statistical software. Demographic variables included chronological age at visit and sex. Clinical assessments included cognitive measures (IQ deviation scores, Stanford-Binet Intelligence Scale, Woodcock-Johnson III achievement test), behavioral assessments (Social Communication Questionnaire [SCQ] total score, Anxiety, Depression, and Mood Scale [ADAMS] subscales for mania, depression, social avoidance, anxiety, and obsessive-compulsive behaviors), and autism-related behaviors (Aberrant Behavior Checklist-Fragile X Syndrome [ABC-FXS] subscales for irritability, lethargy, stereotypy, hyperactivity, speech, and social avoidance).

## Results

In this study, we utilized a previously published resting-state EEG dataset from our laboratory^16^, comparing source-localized recordings between 70 participants with a FXS diagnosis and 71 age- and sex-matched typically developing controls (TDC) (**Fig. 2**, **Table 1**). Neuropsychiatric assessments were used to estimate clinical phenotypic differences between groups. Raw data were blindly preprocessed, myogenic contamination and other artifacts were removed, and the data was filtered to isolate the alpha frequency band (8-12.5 Hz).

**Fig. 2.**
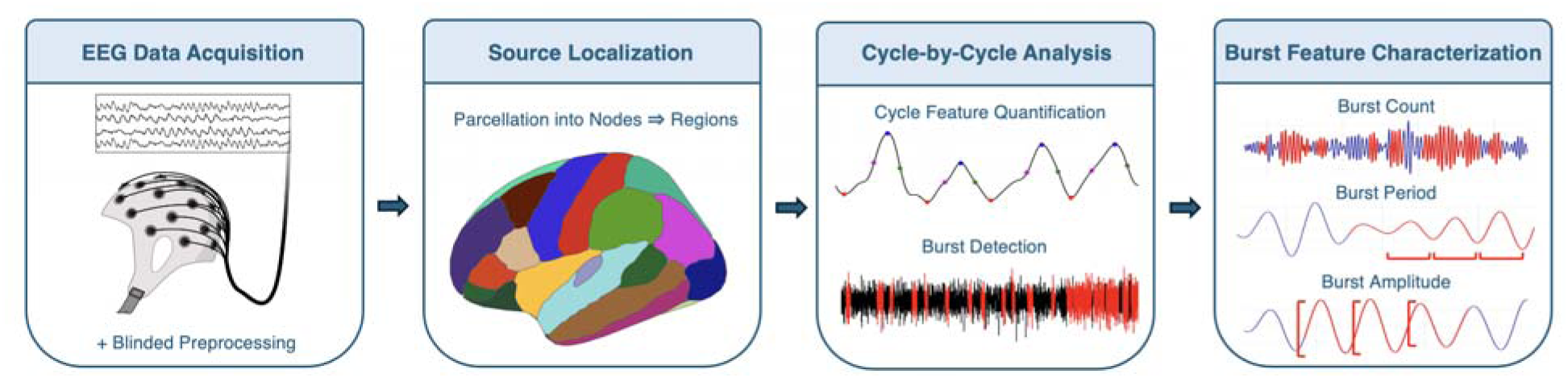
Overview of methodology. The analytical pipeline consists of four sequential stages. **(1) EEG Data Acquisition**: 128-channel resting-state EEG recordings undergo blinded preprocessing including filtering, artifact removal, and channel interpolation. **(2) Source Localization**: Weighted minimum norm estimation (MNE) reconstructs cortical activity, with 15,000 vertices parcellated into 68 Desikan-Killiany atlas nodes and grouped into 14 bilateral cortical regions^65^. **(3) Cycle-by-Cycle Analysis**: Alpha-band filtered signals (8-12.5 Hz) undergo burst detection using optimized *bycycle* parameters to identify sustained oscillatory activity on a cycle-by-cycle basis. **(4) Burst Feature Characterization**: Three key waveform features are extracted from detected bursts: burst count (binary occurrence), burst period (cycle duration), and burst amplitude (peak-to-trough magnitude).

Cycle-by-cycle waveform analysis was performed using the *bycycle* package to identify three alpha burst features of interest: burst count, period, and amplitude^37^. These features were analyzed by generalized linear mixed-effects models that included group (FXS or TDC), region (left and right; occipital, cingulate, parietal, temporal, central, frontal, and prefrontal), and sometimes sex (male or female) as fixed effects, with subject and nested node structure as random effects. To examine whether cycle-by-cycle alpha features might relate to behavioral and cognitive functioning, we conducted exploratory analyses testing associations between burst characteristics and clinical measures within the FXS group.

### Reduced alpha burst count in FXS

Alpha bursts occurred significantly less frequently in FXS participants compared to controls, with an 11% reduction in burst count (TDC/FXS = 1.11, z = 3.19, P = 0.001, n = 141) (**Fig. 3a,b**). The reduction in alpha burst count showed sex-specific patterns. Males with FXS showed 19% fewer bursts than control males (TDC/FXS = 1.19, z = 4.07, P = 4.63 × 10^-5^, n = 79), whereas females with FXS and control females did not differ (TDC/FXS = 1.03, z = 0.65, P = 0.518, n = 62) (**Fig. 3c,d**).

**Fig. 3.**
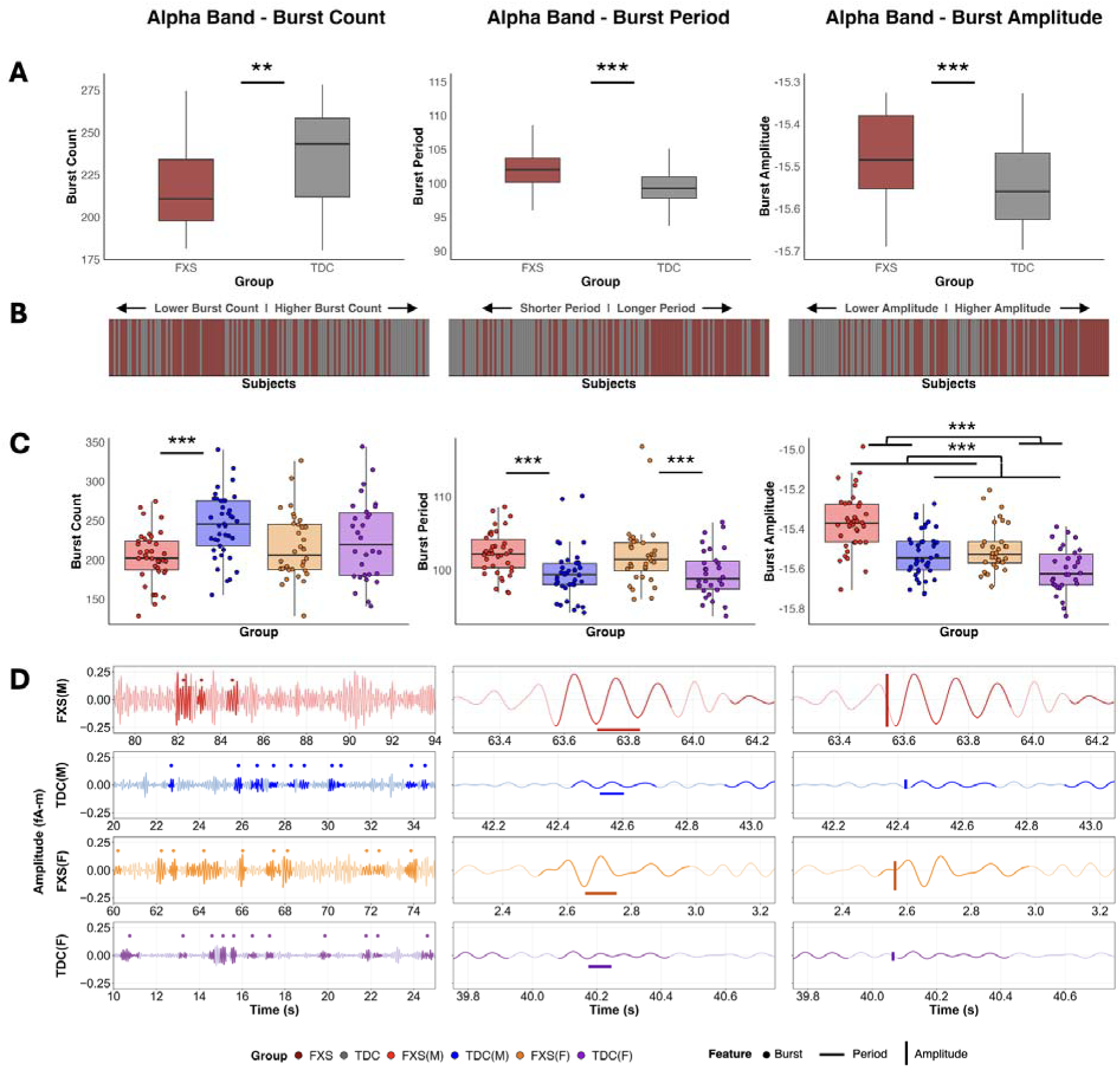
Burst features reveal dysrhythmic alpha burst activity in FXS. **(a)** Group-level comparisons reveal significant differences across all three alpha burst features. FXS participants show reduced burst count (TDC/FXS = 1.11, z = 3.19, P = 0.001, n = 141), prolonged periods indicating slower frequencies (2.92 ms difference; t_9556_ = −4.86, P = 1.20 × 10^-6^), and elevated amplitudes (0.13 log_10_pA-m difference; t_9555_ = −6.23, P = 4.92 × 10^-10^) compared to TDC. **(b)** Subject-level rankings for each burst feature demonstrate the distribution of individual participants along the feature continuum. Bars are color-coded by group (FXS: dark red; TDC: gray). **(c)** Sex-stratified analysis with individual participant data points overlaid on boxplots demonstrates that burst count reductions are driven primarily by FXS males (TDC/FXS = 1.19, z = 4.07, P = 4.63 × 10^-5^, n = 79), while FXS females show no significant difference from TDC females (TDC/FXS = 1.03, z = 0.65, P = 0.518, n = 62). Period prolongation occurs in both FXS sexes, and amplitude elevation shows a main effect of sex with males > females across both groups (0.10 log_10_pA-m; t_9555_ = 5.00, P = 5.94 × 10^-7^). **(d)** Representative alpha-filtered raw time series signals from each group-sex combination showing cycle-by-cycle burst detection across three features: burst count (left), period (middle), and amplitude (right). Bold highlighting and associated filled dots above indicate identified bursts, horizontal bars indicate example burst periods, and vertical bars indicate example burst amplitudes. **P < 0.01; ***P < 0.001.

*Parietal region reductions*. Regionally, alpha burst count was significantly reduced in FXS compared to TDC participants, and widespread across most cortical regions (**Fig. 4a**). The largest reductions were observed in the posterior association regions, notably the parietal cortices bilaterally (RP: TDC/FXS = 1.19, z = 4.38, P = 1.68 × 10^-4^, n = 141; LP: TDC/FXS = 1.16, z = 3.92, P = 6.18 × 10^-4^, n = 141) (**Fig. 4d** and Supplementary Table 2). The frontal and prefrontal cortices were the only regions not significantly different between FXS and TDC groups (**Fig. 4d** and Supplementary Table 2).

**Fig. 4.**
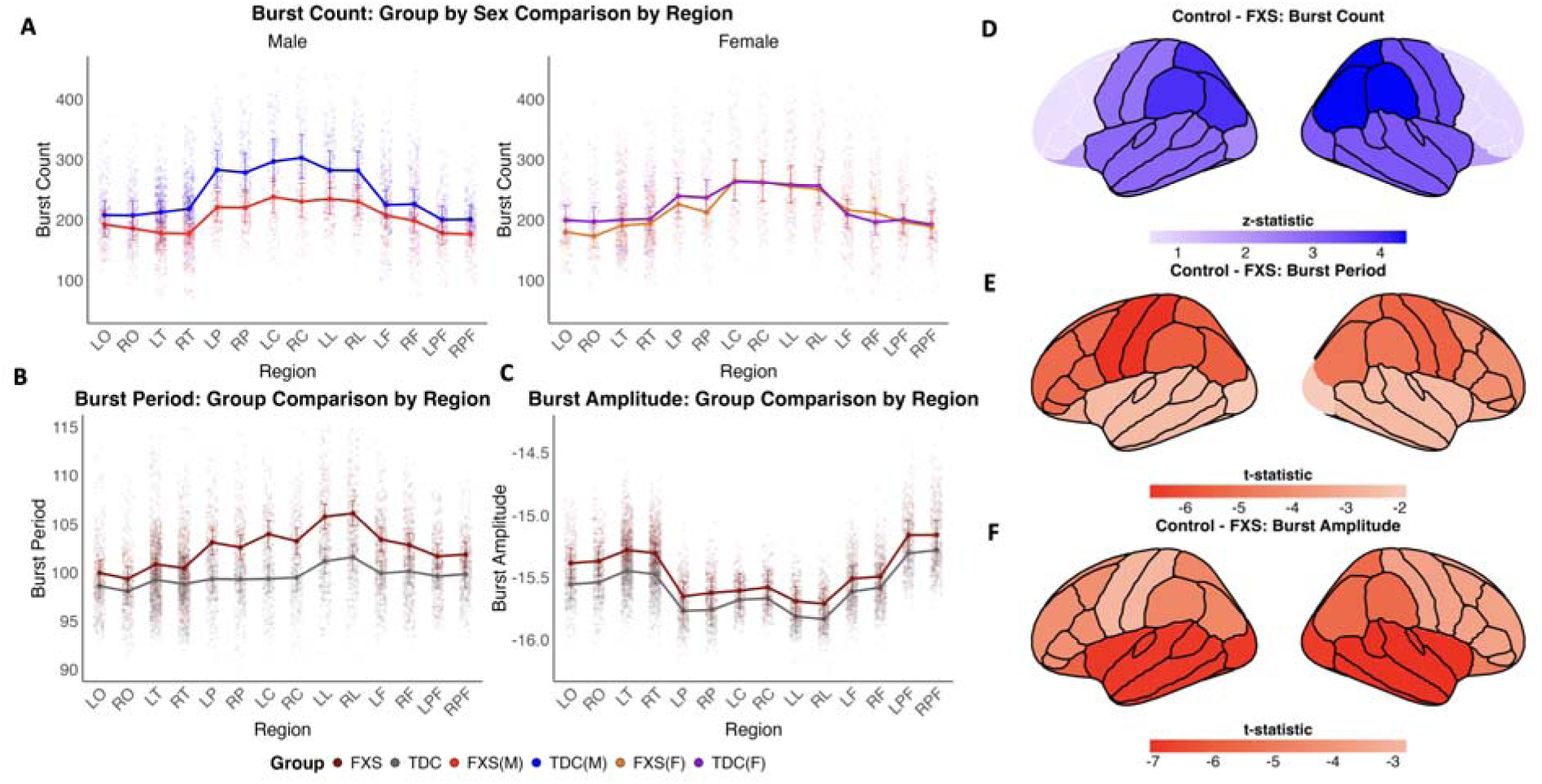
Alpha burst abnormalities demonstrate distinct patterns across cortical areas. **(a)** Group × region × sex interaction for burst count demonstrates that FXS males (red) show widespread reductions in burst occurrence compared to TDC males (blue) across most cortical regions, while FXS females (orange) show no significant differences from TDC females (purple). **(b)** Group × region analysis for burst period reveals prolonged alpha cycles in FXS compared to TDC, with the largest differences observed in central and cingulate regions. **(c)** Amplitude analysis shows elevated alpha burst strength in FXS, with particularly pronounced increases in temporal and sensory regions. Regions are arranged from sensory (left) to cognitive control areas (right). Regions shown as two-letter abbreviations from DK cortical atlas (Supplementary Table 3)^65^. Values on the y-axes are estimated marginal means (emmeans). Error bars represent 95% confidence intervals around the estimated marginal means. **(d-f)** Cortical surface maps based on the DK cortical atlas^64^ (Supplementary Fig. 1) display statistical contrasts (t-values) for TDC - FXS (negative values (red) indicate FXS > TDC; positive vales (blue) indicate TDC > FXS). Black outlines indicate regions with significant group differences (P < 0.05). **(d)** Burst count reductions (blue regions) are most prominent in parietal cortices (RP: TDC/FXS = 1.19, *z* = 4.38, P = 1.68 × 10^-4^, n = 141; LP: TDC/FXS = 1.16, *z* = 3.92, P = 6.18 × 10^-4^, n = 141). **(e)** Period prolongation (red regions) shows strongest effects in cingulate and central areas: left cingulate (LL: 4.6 ms; t_9556_ = −6.74, P = 1.52 × 10^-10^), right cingulate (RL: 4.5 ms; t_9556_ = −6.64, P = 1.52 × 10^-10^), left central (LC: 4.6 ms; t_9556_ = −6.65, P = 1.52 × 10^-10^), and right central (RC: 3.8 ms; t_9556_ = −5.41, P = 1.78 × 10^-7^). **(f)** Burst amplitude elevation in FXS (red regions) is most pronounced in sensory cortices: right temporal (RT: 0.17 log_10_pA-m; t_9555_ = - 7.04, P = 2.78 × 10^-11^), left temporal (LT: 0.17 log_10_pA-m; t_9555_ = −6.88, P = 3.06 × 10^-11^), right occipital (RO: 0.17 log_10_pA-m; t_9555_ = −6.83, P = 3.15 × 10^-11^), and left occipital (LO: 0.17 log_10_pA-m; t_9555_ = −6.88, P = 3.06 × 10^-11^).

*Positive age association.* Whole-brain alpha burst count showed a significant positive correlation with age (adjusted R² = 0.130, f² = 0.150, P = 0.023, n = 69), with this relationship being significant in females (adjusted R² = 0.138, f² = 0.161, P = 0.022, n = 31) but only trending in males (adjusted R² = 0.054, f² = 0.057, P = 0.087, n = 38), suggesting sex-specific developmental changes in alpha burst frequency across the lifespan in FXS (**Fig. 5e,f**).

**Fig. 5.**
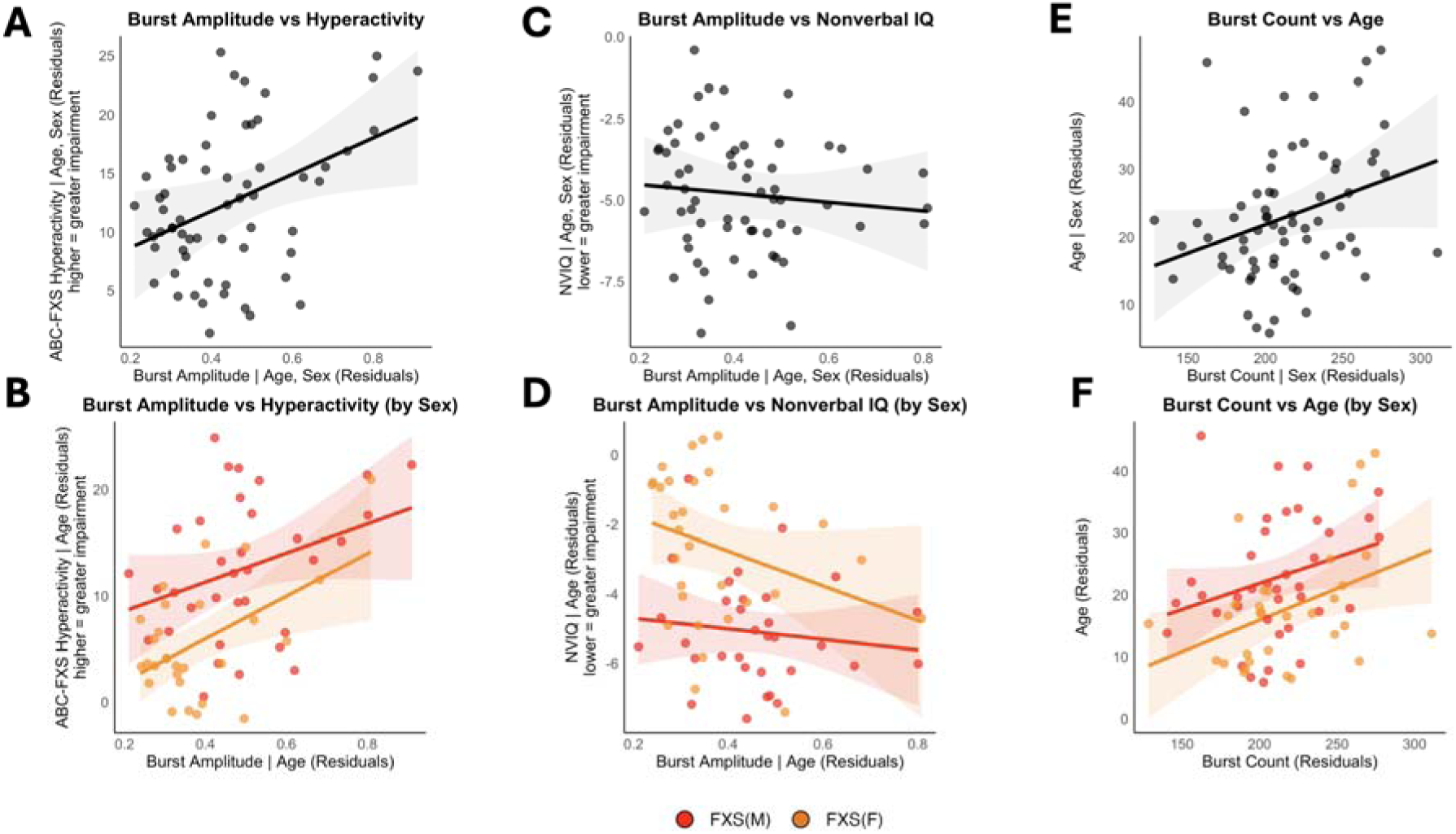
**Alpha burst amplitude scales with hyperactivity and cognitive impairment in FXS**. The top row shows interaction models examining sex differences in the relationships between neural oscillation parameters and clinical outcomes. The bottom row displays the same relationships stratified by sex (orange = female, red = male). **(a-b)** Higher alpha burst amplitude was associated with greater hyperactivity scores on the ABC-FXS scale (adj R² = 0.431, f² = 0.756, P = 0.029, n = 62), with this relationship driven primarily by females (adj R² = 0.285, f² = 0.399, P = 0.008, n = 28) compared to males (adj R² = 0.335, f² = 0.505, P = 0.088, n = 34). **(c-d)** Lower nonverbal IQ scores showed a trending association with higher burst amplitude (adj R² = 0.393, f² = 0.649, P = 0.051, n = 63), with no significant sex interaction (P = 0.281). When analyzed separately by sex, neither males (adj R² = 0.183, f² = 0.225, P = 0.495, n = 32) nor females (adj R² = 0.079, f² = 0.086, P = 0.111, n = 31) showed significant associations. **(e-f)** Alpha burst count was positively associated with age (adj R² = 0.130, f² = 0.150, P = 0.023, n = 69), with no significant sex interaction (P = 0.786). This relationship was significant in females (adj R² = 0.138, f² = 0.161, P = 0.022, n = 31) but only marginal in males (adj R² = 0.054, f² = 0.057, P = 0.087, n = 38). All models controlled for age except the burst count-age relationship, which controlled for sex. Gray ribbons and shaded areas represent 95% confidence intervals. Points represent individual participants with residual values after controlling for covariates.

### Increased alpha burst period in FXS

When alpha bursts did occur, burst periods were significantly prolonged by approximately 3 ms in the FXS group (t_9556_ = −4.86, P = 1.20 × 10^-6^) (**Fig. 3a,b**), and there were no sex effects or group by sex interactions observed (**Fig. 3c,d**).

*Regional gradients in prolongation.* Alpha burst period was significantly longer in FXS across most cortical regions, with the strongest effects in cingulate and central cortices bilaterally, showing an approximate 4 ms prolongation: left cingulate (LL: 4.6 ms; t_9556_ = −6.74, P = 1.52 × 10^-10^), right cingulate (RL: 4.5 ms; t_9556_ = −6.64, P = 1.52 × 10^-10^), left central (LC: 4.6 ms; t_9556_ = - 6.65, P = 1.52 × 10^-10^), and right central (RC: 3.8 ms; t_9556_ = −5.41, P = 1.78 × 10^-7^) (**Fig. 4b,e** and Supplementary Table 2). In contrast, this prolongation diminished progressively toward sensory cortices, where occipital and temporal regions showed substantially smaller period changes: right occipital (RO: 1.3 ms; t_9556_ = −1.88, P = 0.060), left occipital (LO: 1.4 ms; t_9556_ = - 1.99, P = 0.050), right temporal (RT: 1.6 ms; t_9556_ = −2.49, P = 0.016), and left temporal (LT: 1.6 ms; t_9556_ = −2.46, P = 0.016) (**Fig. 4b,e** and Supplementary Table 2).

*No clinical associations*. Whole-brain burst period was not significantly associated with any clinical measures.

### Elevated alpha burst amplitudes in FXS

Burst amplitudes were elevated in FXS (0.13 log_10_pA-m difference; t_9555_ = −6.23, P = 4.92 × 10^-10^), indicating stronger individual oscillatory pulses (**Fig. 3a,b**). A main effect of sex was observed, with males showing higher alpha burst amplitude than females overall (0.10 log_10_pA-m; t_9555_ = 5.00, P = 5.94 × 10^-7^), and both FXS males and females exhibited elevated amplitudes relative to their sex-matched controls (**Fig. 3c,d**). There was no group by sex interaction detected.

*Sensory region elevations.* While alpha burst amplitude was significantly elevated in FXS across all cortical regions (**Fig. 4c,f**), there were particularly strong increases in sensory-oriented regions, including temporal and occipital cortices bilaterally: right temporal (RT: 0.17 log_10_pA-m; t_9555_ = −7.04, P = 2.78 × 10^-11^), left temporal (LT: 0.17 log_10_pA-m; t_9555_ = −6.88, P = 3.06 × 10^-11^),right occipital (RO: 0.17 log_10_pA-m; t_9555_ = −6.83, P = 3.15 × 10^-11^), and left occipital (LO: 0.17 log_10_pA-m; t_9555_ = −6.88, P = 3.06 × 10^-11^) (**Fig. 4c,f** and Supplementary Table 2), precisely where period prolongation was minimal. Central regions, which showed especially increased burst period, exhibited the smallest amplitude elevations (LC: 0.071 log_10_pA-m; t_9555_ = −2.78, P = 0.005; RC: 0.089 log_10_pA-m; t_9555_ = −3.51, P = 4.92 × 10^-4^) (**Fig. 4c,f** and Supplementary Table 2).

*Positive hyperactivity association.* Higher whole-brain burst amplitudes were significantly associated with increased hyperactivity scores (adjusted R² = 0.431, f² = 0.756, P = 0.029, n = 62), with this relationship being particularly pronounced in females (adjusted R² = 0.285, f² = 0.399, P = 0.008, n = 28) compared to males (adjusted R² = 0.335, f² = 0.505, p = 0.088, n = 34) (**Fig. 5a,b**). Burst amplitude showed a trending negative association with general intelligence (NVIQ) scores (adjusted R² = 0.393, f² = 0.649, P = 0.051, n = 63), indicating that higher amplitude bursts may correspond to poorer cognitive performance, though sex-stratified analyses revealed no significant associations in either males or females (**Fig. 5c,d**).

### Cross-feature spatial dissociation of burst abnormalities across cortical regions

Alpha bursts in FXS are characterized by fewer but stronger and slower oscillatory events that do not affect all cortical regions uniformly. Regions exhibiting the strongest prolongation (increased burst period), notably cingulate and central cortices, showed minimal amplitude elevations, whereas sensory cortices, temporal and occipital, displaying the largest amplitude increases demonstrated less time-related change (**Fig. 4**). Burst count reductions aligned most closely with regions showing prolonged period rather than elevated amplitude. Therefore, regions characterized by complex inter-regional connectivity showed predominant time-associated dysregulation, whereas primarily sensory cortices showed amplitude-dominant patterns, potentially reflecting distinct circuit-level properties.

## Discussion

Deviations in alpha oscillatory power are frequently cited across various neuropsychiatric conditions from Parkinson’s disease to ASD^8, 10^, however, this lack of specificity has also limited its utility as a biomarker. Alpha oscillations’ central role in coordinating cortical networks and their vulnerability across neuropathologies^48, 49^ suggests that more mechanistic characteristics of alpha dynamics beyond aggregate power may hold promise as prognostic or therapeutic markers. Conventional power spectral analysis collapses time and, in relative metrics, normalizes by total broadband activity, thereby obscuring burst structure and permitting the coexistence of higher absolute yet lower relative alpha power.

FXS, an X-linked monogenic disorder, is associated with pronounced and replicable sex-specific alterations in alpha spectral profiles among neurodevelopmental conditions^16, 18, 50^. FXS arises from a near-complete loss of a single protein (FMRP) with well-characterized disruption of synaptic and circuit function^21, 23, 24^. It therefore represents a critical model for testing whether mechanistic features of alpha dynamics can be linked to clinical phenotypes and predictions of circuit level dysfunction. We conducted a discontinuous cycle-by-cycle burst analysis, decomposing aggregate power into distinct burst features, providing more specific targets. This approach revealed that alpha activity in FXS is not simply stronger or weaker, but fundamentally dysrhythmic.

We found that FXS is characterized by alpha bursts that have (1) reduced burst count specifically in males, (2) prolonged burst periods across both sexes, (3) increased burst amplitudes in both sexes, with males showing the most pronounced increases, and (4) spatially dissociated patterns, with burst count and burst period alterations greatest in posterior-parietal and midline-central regions, while burst amplitude increases were maximal in sensory cortices. These findings delineate the time-associated structure of alpha dysregulation in FXS, identifying burst-level signatures that enable hypothesis generation about thalamocortical and interneuron mechanisms and their potential as translational biomarkers.

### Dysrhythmic alpha burst dynamics address paradoxical power findings

Alpha bursts in FXS are less frequent, slower, yet higher amplitude relative to controls, as reflected by decreased burst count, prolonged burst periods, and increased burst amplitude. Since spectral power scales with the square of amplitude, increased burst amplitude can disproportionately elevate absolute alpha power^51^. When paired with reduced burst count and longer burst period, the time-related density of alpha activity is decreased, yielding a lower relative power when normalized by total broadband activity, consistent with our previous work^16^. *Bycycle* quantifies burst-level timing and morphology in the time domain; while burst amplitude contributes to absolute power, this method captures time-related density and waveform shape that are not represented by absolute or relative power estimates. Furthermore, this burst-level analytic framework may be applicable to other neurodevelopmental and neuropsychiatric disorders characterized by oscillatory dysrhythmias, enabling cross-condition comparisons and refining circuit-level biomarkers.

### Alpha burst features demonstrate complex sex effects

Sex-specific patterns in alpha oscillatory dysfunction appear to align with established phenotypic differences between males and females with FXS^25^. The reduction in alpha burst count was driven entirely by males, with FXS females showing burst counts statistically indistinguishable from control females. This preservation may suggest that FMRP expression from random X-inactivation in females is sufficient to maintain burst generation. Burst amplitude showed elevations in both FXS males and females compared to same-sex controls, with FXS males exhibiting the highest values overall. This pattern likely reflects both FMRP-dependent processes as well as baseline sex differences, as even TDC males had higher burst amplitudes than TDC females. Alpha period prolongation occurred across both sexes in the FXS group, representing a fundamental aspect of FXS pathophysiology, as even millisecond-scale period shifts can potentially disrupt spike-timing precision and cross-regional phase coherence^20, 52^.

These mechanistic deviations likely contribute to impaired temporal coordination of neural activity, which has been directly linked to core behavioral deficits in FXS, including sensory hypersensitivity, attention instability, and social-communication impairments^16, 53–55^. This finding of sex-independent period prolongation in FXS has important implications for biomarker development. While previous FXS research has focused predominantly on males due to their more severe phenotypic presentation^56^, alpha period provides a quantifiable neural signature that is equally detectable in both sexes, enabling more inclusive approaches to future clinical trials and mechanistic studies.

### Dissociable alpha rhythms: gradient regional specificity informs mechanisms

Our regional analysis reveals that while alpha dysregulation occurs mostly globally in FXS, distinct properties of the alpha rhythm show their strongest effects in specific cortical areas, suggesting spatially dissociable alpha dysfunction. Timing-associated dysregulation, prolonged periods and reduced burst counts, was most pronounced in posterior association cortices (parietal) and cognitive control regions (central, cingulate), areas serving as hubs within attentional and executive networks^57^. In contrast, amplitude elevations were strongest in sensory cortices (occipital, temporal), regions where alpha is thought to gate sensory processing through inhibitory mechanisms^58^. This graded pattern of regional differences suggests that the widespread oscillatory dysfunction in FXS, stemming from the absence of FMRP, interacts with the unique functional demands and circuit properties of different brain regions. This spatial dissociation may further reflect differential vulnerability of alpha generation mechanisms across the cortical hierarchy. Cortical regions generate alpha oscillations that propagate across cortical areas in hierarchical patterns from higher-order to lower-order regions^6^. Our findings suggest that the timing-related dominant patterns in higher-order regions may reflect disrupted network-level coordination, and the amplitude-dominant patterns in lower-order sensory cortices may reflect local circuit properties of potentially enhanced pyramidal neuron synchrony. Critically, these patterns emerged from resting-state recordings without task demands, indicating they reflect intrinsic circuit dysfunction rather than state-dependent engagement.

### Alpha burst characteristics associated with clinical measures in FXS

The present exploratory findings reveal complex relationships between alpha burst characteristics and clinical phenotypes in FXS, with notable sex-specific patterns. While our previous work demonstrated that reduced relative alpha power correlated with anxiety, auditory attention, and social functioning deficits^16^, burst amplitude emerged as the most clinically relevant feature, showing significant associations with hyperactivity behaviors and trending associations with general intelligence. This divergence suggests that different aspects of alpha dynamics, aggregate relative spectral power compared to burst-level characteristics, may relate to distinct symptom domains in FXS. Hyperactivity is an established characteristic of FXS, notable in both humans and associated *Fmr1* KO mice^25, 59^. The relationship with hyperactivity was particularly pronounced in females, suggesting possible sex-dependent neural mechanisms underlying behavioral regulation in FXS. The association between higher burst amplitudes and lower generalized IQ, while not reaching conventional significance, suggests potential relationships between oscillatory strength and cognitive performance that warrant further investigation. The developmental trajectory of burst count, showing positive associations with age particularly in females, aligns with previous findings of increased PAF with development^60, 61^, potentially indicating that alpha burst generation may reflect maturational processes. These findings position alpha burst amplitude as a potentially valuable neurobiological marker that scales with core behavioral phenotypes, though the mechanisms underlying these relationships require further investigation.

### Interneuron circuit dysfunction as unifying mechanism

The distinct cycle-by-cycle features of alpha dysregulation in FXS, reduced burst count, prolonged periods, and increased amplitudes, are consistent with interneuron circuit dysfunction, particularly involving parvalbumin (PV) and somatostatin (SOM) expressing GABAergic interneurons. Recent computational work by Tahvili et al. demonstrates that PV and SOM interneurons play distinct causal roles in controlling network oscillations, with SOM cells primarily modulating oscillation amplitude while PV cells regulate frequency and network stability^62^. The CAMINOS (canonical microcircuit network oscillations) model reveals that increasing SOM-to-PV ratios produces oscillatory dynamics at lower frequencies, transitioning from gamma frequencies toward slower alpha/beta oscillations.

There is a well-established reduction of neocortical PV+ interneurons in FXS^63, 64^. Evidence for SOM interneuron alterations is relatively unexplored in FXS, although a preservation of SOM density has been reported^65^. However, prolonged loss of PV cells may trigger increased SOM interneuron activity to compensate for altered excitation-inhibition balance, representing homeostatic circuit stabilization^66, 67^. If PV interneurons are reduced while SOM interneurons are relatively preserved or increased, the effective SOM/PV ratio would increase. Based on the computational framework, this altered ratio could explain our key findings: decreased PV function may account for prolonged cycle periods and reduced burst occurrence (reflecting decreased frequencies and network stability), while relatively increased SOM influence could explain the increased amplitudes observed across both sexes (**Fig. 6**).

**Fig. 6.**
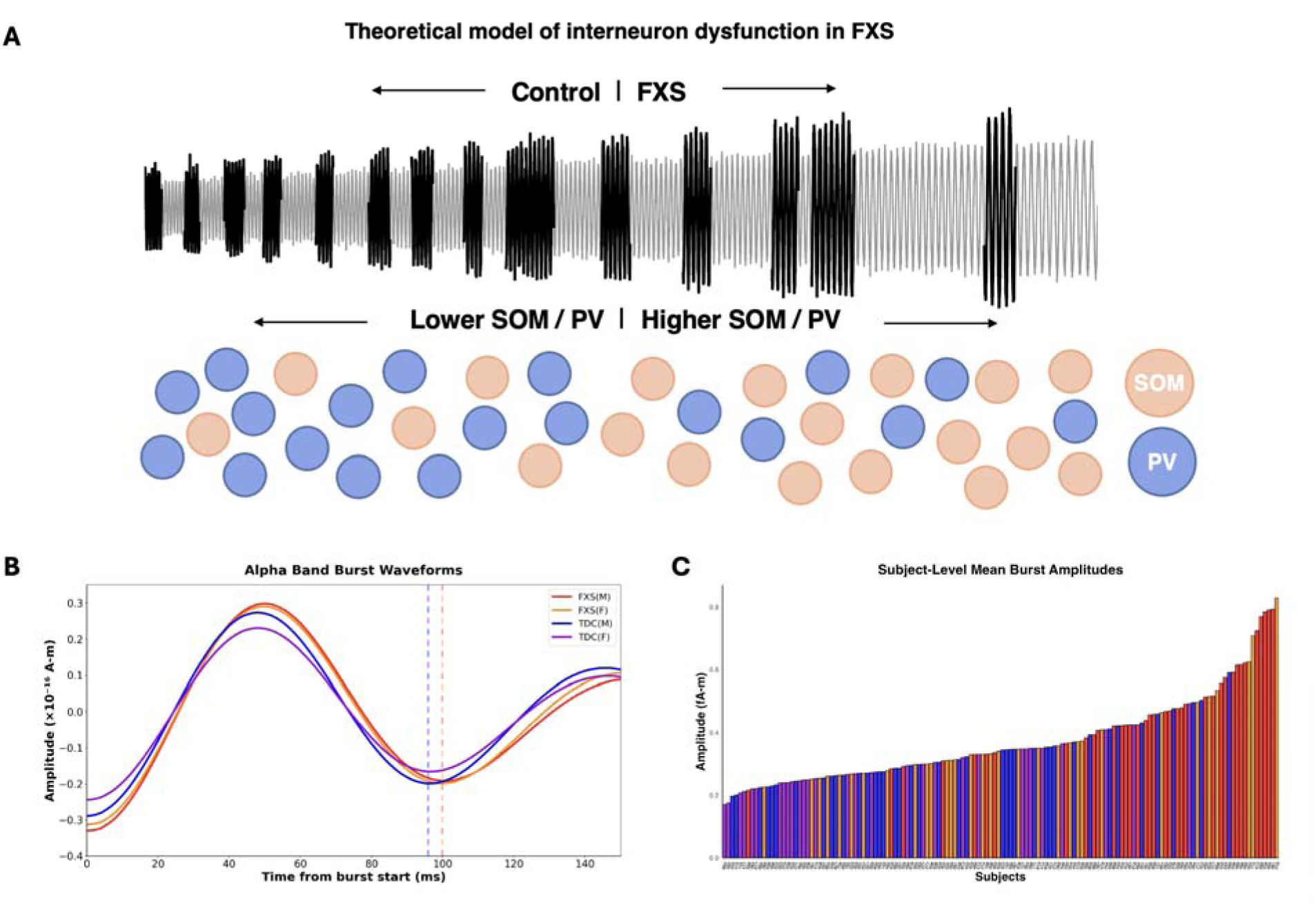
Model of hypothesized interneuron circuit dysfunction underlying alpha oscillation dysregulation in FXS. **(a)** Simulated alpha oscillations (grey) demonstrate how increasing SOM/PV interneuron ratios (left to right) reproduce the characteristic transition from TDC to FXS oscillatory patterns. Black regions indicate alpha bursts. Orange circles represent PV interneurons, which decrease in density from left to right. Blue circles represent SOM interneurons, with SOM/PV ratios progressively increasing. As the SOM/PV ratio increases, alpha oscillations exhibit the hallmark FXS features: reduced burst frequency, prolonged burst periods, and increased burst amplitudes. This computational framework suggests that PV interneuron loss combined with relative SOM interneuron preservation drives the distinct cycle-by-cycle alpha dysregulation observed in FXS, with reduced PV function accounting for decreased network stability and prolonged periods, while increased SOM influence contributes to enhanced oscillation amplitudes. **(b)** Average waveforms of alpha bursts for each group: FXS males (red), FXS females (orange), TDC males (blue), and TDC females (purple). Waveforms aligned by the first trough (time at 0) and axes limited to emphasize first cycle differences (0-150 ms). Vertical dashed lines indicate period of the first cycle within the burst. **(c)** Ranked bar plot showing average alpha band burst amplitudes for individual subjects, ordered from lowest to highest amplitude. Each bar represents one subject’s mean burst amplitude with colors indicating group membership: FXS males (red), TDC males (blue), FXS females (orange), and TDC females (purple). Analysis excludes one outlier subject (241) with amplitude exceeding 3 standard deviations above the group mean.

### Cross-species translation and mechanistic validation

A critical next step involves validating these findings in *Fmr1* KO mouse models to address whether the observed alpha burst feature alterations represent compensatory mechanisms or primary pathological processes. While translational studies have established consistent gamma abnormalities across species, alpha oscillations present a more complex picture. Resting-state studies in *Fmr1* KO mice show conflicting results, with some demonstrating reduced relative alpha power similar to human FXS while others show no differences from controls^31, 32^. Future studies should apply cycle-by-cycle analysis to resting-state recordings in *Fmr1* KO mice to determine whether the specific burst characteristics we identified are conserved across species. Further, this *bycycle* approach will allow us to explore the conflicting results surrounding alpha power in *Fmr1* KO mice, investigating the contributions of reduced alpha burst count and increased burst amplitude that may be concealing differences in alpha power. Crucially, these studies should incorporate quantification of SOM/PV interneuron ratios across brain regions through immunohistochemistry or optogenetic approaches, enabling direct testing of our proposed mechanistic framework. Selective silencing or activation of PV versus SOM interneuron populations could establish causal relationships between altered interneuron balance and specific cycle-by-cycle features, addressing the fundamental question of whether these oscillatory changes drive clinical symptoms or represent compensatory responses.

### Biomarker potential and therapeutic applications

Our cycle-by-cycle alpha features represent valuable biomarkers with immediate therapeutic potential. The quantifiable nature of burst count, period, and amplitude provides objective metrics that could enhance sensitivity in clinical trials, better stratify patient groups, and guide treatment development. For BCI applications, real-time detection of these aberrant alpha dynamics could trigger targeted neuromodulation to restore healthy oscillatory patterns^68^. The correlation between alpha amplitude and symptom severity suggests these metrics could serve as both treatment targets and outcome measures, providing a neurobiologically-grounded approach to monitoring therapeutic progress that complements traditional behavioral assessments.

### Limitations

Since our data was collected during resting-state, to test hypotheses related to how alpha dysfunction in FXS relates to region-specific function, future studies will need to explore how alpha burst features are modulated during cognitive and sensory task-based paradigms. For example, we would predict that during auditory stimulation, burst amplitude specifically will increase in the temporal regions. Further, this study is limited by its cross-sectional design, which precludes assessment of longitudinal changes in alpha burst dynamics. Additionally, while the effect of non-epileptic medications on our results cannot be disregarded, previous EEG studies in neuropsychiatric conditions have shown minimal associations with medication variables^69^, including in FXS^53^. Further, while source localization remains limited by the inverse problem, it is currently the best method to achieve cortical spatial resolution^40^ in populations with sensory hypersensitivities such as in FXS, where MRI-based methods are often not feasible.

From a methodological perspective, no single cycle feature fully characterizes waveform physiology, so our focus on three specific metrics (burst count, period, and amplitude) may not capture all relevant aspects of oscillatory dysfunction^37^. These features were selected to disambiguate the paradoxical alpha power findings, but additional analyses looking at state transition dynamics or within-burst feature modulation may reveal a more complete picture of alpha dysregulation in FXS. This study is a post hoc analysis, so such exploratory findings would benefit from replication in future cohorts. Lastly, FMRP expression was not quantified for participants, limiting potential stratification by molecular severity as it may contribute to within-group variability.

### Conclusions

This cycle-by-cycle analysis reveals that FXS is characterized by complex alpha oscillatory dysfunction that extends beyond simple power reductions to encompass fundamental alterations in burst dynamics and temporal structure. While alpha oscillations remain one of the most robust yet mechanistically underspecified signals in clinical neuroscience, this work demonstrates that moving beyond aggregate power to cycle-by-cycle burst features reveals disorder-specific, sexually dimorphic, and regionally selective signatures that traditional spectral approaches obscure. These findings demonstrate that different dimensions of alpha physiology (burst count, period, and amplitude) may fractionate along distinct symptom domains and disorders, offering a path toward objective, mechanistically grounded biomarkers. Alpha dysregulation represents a final common pathway whose precise parameterization may ultimately guide circuit-based stratification and individualized therapeutic strategies across neurodevelopmental and neuropsychiatric disorders.

## Declarations

### Ethics approval and consent to participate

All procedures were conducted in accordance with the Declaration of Helsinki. All participants provided written informed consent (or assent, where applicable), and the study was approved by the institutional review board of Cincinnati Children’s Hospital Medical Center.

## Consent for publication

Not applicable.

## Availability of data and materials

The de-identified datasets and code used for analysis in this study are available from the corresponding author upon request. EEG datasets are available at https://doi.org/10.5281/zenodo.6385768. Correspondence and material requests should be addressed to Peyton Siekierski.

## Acknowledgements

We thank the participants and families involved at the Cincinnati Fragile X Research and Treatment Center who participated in this study.

## Authors’ Contributions

P.S. contributed to study conceptualization, data analysis, figure generation, and manuscript writing. Y.L. and G.W. contributed to data collection, original study conceptualization, and manuscript feedback. R.E. and D.G. provided manuscript feedback. Z.E. and C.A.E. contributed to study conceptualization. E.V.P. contributed to study conceptualization, supervision, manuscript editing, and interpretation of results. All authors reviewed and approved the final manuscript.

## Abbreviations

ABC: Aberrant Behavior Checklist
ADAMS: Anxiety, Depression, and Mood Scale
DK: Desikan-Killiany (atlas)
EEG: electroencephalography
EGI: Electrical Geodesics, Inc.
FMRP: fragile X messenger ribonucleoprotein
FXS: Fragile X syndrome
GLMM: generalized linear mixed-effects models
IQ: intelligence quotient
KO: knockout
LC: Left Central
LF: Left Frontal
LL: Left Cingulate
LO: Left Occipital
LP: Left Parietal
LPF: Left Prefrontal
LT: Left Temporal
MNE: minimum norm estimate
MRI: magnetic resonance imaging
NVIQ: nonverbal intelligence quotient
PAF: peak alpha frequency
PV: parvalbumin
RC: Right Central
RF: Right Frontal
RL: Right Cingulate
RO: Right Occipital
RP: Right Parietal
RPF: Right Prefrontal
RT: Right Temporal
SCQ: Social Communication Questionnaire
SOM: somatostatin
TDC: typically developing controls
ABC: Aberrant Behavior Checklist
ADAMS: Anxiety, Depression, and Mood Scale
DK: Desikan-Killiany (atlas)
EEG: electroencephalography
EGI: Electrical Geodesics, Inc.
FMRP: fragile X messenger ribonucleoprotein
FXS: Fragile X syndrome
GLMM: generalized linear mixed-effects models
IQ: intelligence quotient
KO: knockout
LC: Left Central
LF: Left Frontal
LL: Left Cingulate
LO: Left Occipital
LP: Left Parietal
LPF: Left Prefrontal
LT: Left Temporal
MNE: minimum norm estimate
MRI: magnetic resonance imaging
NVIQ: nonverbal intelligence quotient
PAF: peak alpha frequency
PV: parvalbumin
RC: Right Central
RF: Right Frontal
RL: Right Cingulate
RO: Right Occipital
RP: Right Parietal
RPF: Right Prefrontal
RT: Right Temporal
SCQ: Social Communication Questionnaire
SOM: somatostatin
TDC: typically developing controls

