## Supplemental Materials for "Alpha oscillations are dysrhythmic in Fragile X syndrome"

### Supplementary Materials

| **Feature** | **Model Formula** |
| --- | --- |
| Burst Count | group × sex × region + (1\|eegid) + (1\|eegid:region) + (1\|region:node) |
| Period | group × region + (1\|eegid) + (1\|eegid:region) + (1\|region:node) |
| Amplitude | group × region + sex + (1\|eegid) + (1\|eegid:region) + (1\|region:node) |

**Supplementary Table 1. Model selection summary.** Models were built using a forward stepwise approach for each alpha burst feature. The modeling process began with group as a fixed effect, then sequentially evaluated the addition of sex, region, and their interactions (group × sex, group × region, etc.). This iterative approach continued until no further significant effects or interactions could be identified, resulting in the final model specifications above.

| **Region** | **Amplitude Δ (log₁₀pA-m)** | **Amplitude Statistics** | **Amplitude P (FDR)** | **Period Δ (ms)** | **Period Statistics** | **Period P (FDR)** | **Burst Count Ratio (TDC/FXS)** | **Burst Count Statistics** | **Burst Count P (FDR)** |
| --- | --- | --- | --- | --- | --- | --- | --- | --- | --- |
| Left Occipital | -0.173 ± 0.025 | t(9555) =  -6.88 | 3.06e-11*** | -1.36 ± 0.68 | t(9556) =  -1.99 | 0.0498* | 1.10 | z = 2.34 | 0.0270* |
| Right Occipital | -0.171 ± 0.025 | t(9555) =  -6.83 | 3.15e-11*** | -1.29 ± 0.68 | t(9556) =  -1.88 | 0.0601 | 1.13 | z = 3.09 | 0.0057** |
| Left Temporal | -0.167 ± 0.024 | t(9555) =  -6.88 | 3.06e-11*** | -1.63 ± 0.66 | t(9556) =  -2.46 | 0.0160* | 1.12 | z = 2.90 | 0.0088** |
| Right Temporal | -0.171 ± 0.024 | t(9555) =  -7.04 | 2.78e-11*** | -1.64 ± 0.66 | t(9556) =  -2.49 | 0.0160* | 1.13 | z = 3.18 | 0.0052** |
| Left Parietal | -0.114 ± 0.025 | t(9555) =  -4.53 | 8.35e-06*** | -3.80 ± 0.68 | t(9556) =  -5.55 | 1.04e-07*** | 1.16 | z = 3.92 | 6.18e-04*** |
| Right Parietal | -0.138 ± 0.025 | t(9555) =  -5.49 | 9.57e-08*** | -3.31 ± 0.68 | t(9556) =  -4.84 | 2.68e-06*** | 1.19 | z = 4.38 | 1.68e-04*** |
| Left Central | -0.071 ± 0.026 | t(9555) =  -2.78 | 0.0054** | -4.63 ± 0.70 | t(9556) =  -6.65 | 1.52e-10*** | 1.11 | z = 2.71 | 0.0118* |
| Right Central | -0.089 ± 0.026 | t(9555) =  -3.51 | 4.92e-04*** | -3.77 ± 0.70 | t(9556) =  -5.41 | 1.78e-07*** | 1.14 | z = 3.44 | 0.0027** |
| Left Cingulate | -0.121 ± 0.025 | t(9555) =  -4.83 | 2.18e-06*** | -4.61 ± 0.68 | t(9556) =  -6.74 | 1.52e-10*** | 1.10 | z = 2.48 | 0.0203* |
| Right Cingulate | -0.124 ± 0.025 | t(9555) =  -4.93 | 1.71e-06*** | -4.54 ± 0.68 | t(9556) =  -6.64 | 1.52e-10*** | 1.12 | z = 2.83 | 0.0093** |
| Left Frontal | -0.104 ± 0.025 | t(9555) =  -4.21 | 3.29e-05*** | -3.50 ± 0.68 | t(9556) =  -5.17 | 5.53e-07*** | 1.02 | z = 0.59 | 0.5550 |
| Right Frontal | -0.087 ± 0.025 | t(9555) =  -3.53 | 4.92e-04*** | -2.71 ± 0.68 | t(9556) =  -4.00 | 1.11e-04*** | 1.03 | z = 0.69 | 0.5246 |
| Left Prefrontal | -0.146 ± 0.025 | t(9555) =  -5.82 | 1.66e-08*** | -2.09 ± 0.68 | t(9556) =  -3.05 | 0.0036** | 1.07 | z = 1.75 | 0.0939 |
| Right Prefrontal | -0.121 ± 0.025 | t(9555) =  -4.84 | 2.18e-06*** | -2.04 ± 0.68 | t(9556) =  -2.98 | 0.0040** | 1.08 | z = 1.88 | 0.0757 |

**Supplementary Table 2. Estimated marginal mean differences in alpha burst features per cortical region.** All values represent estimated marginal mean (emmeans) differences between TDC and FXS (TDC - FXS) groups with 5% FDR-corrected p-values. Amplitude values are log_10_-transformed source-localized estimates in picoampere-meters (pA-m). Burst count shows z-values for TDC/FXS ratio (positive z indicates TDC > FXS). *** P < 0.001; ** P < 0.01; * P < 0.05

| **Abbreviation** | **Region Full Name** | **DK Atlas Nodes** |
| --- | --- | --- |
| LO | Left Occipital | cuneusL; lateraloccipitalL; lingualL; pericalcarineL |
| RO | Right Occipital | cuneusR; lateraloccipitalR; lingualR; pericalcarineR |
| LT | Left Temporal | banksstsL; entorhinalL; fusiformL; inferiortemporalL; insulaL; middletemporalL; parahippocampalL; superiortemporalL; temporalpoleL; transversetemporalL |
| RT | Right Temporal | banksstsR; entorhinalR; fusiformR; inferiortemporalR; insulaR; middletemporalR; parahippocampalR; superiortemporalR; temporalpoleR; transversetemporalR |
| LP | Left Parietal | inferiorparietalL; precuneusL; superiorparietalL; supramarginalL |
| RP | Right Parietal | inferiorparietalR; precuneusR; superiorparietalR; supramarginalR |
| LC | Left Central | paracentralL; postcentralL; precentralL |
| RC | Right Central | paracentralR; postcentralR; precentralR |
| LL | Left Cingulate | caudalanteriorcingulateL; isthmuscingulateL; posteriorcingulateL; rostralanteriorcingulateL |
| RL | Right Cingulate | caudalanteriorcingulateR; isthmuscingulateR; posteriorcingulateR; rostralanteriorcingulateR |
| LF | Left Frontal | caudalmiddlefrontalL; parsopercularisL; parstriangularisL; rostralmiddlefrontalL; superiorfrontalL |
| RF | Right Frontal | caudalmiddlefrontalR; parsopercularisR; parstriangularisR; rostralmiddlefrontalR; superiorfrontalR |
| LPF | Left Prefrontal | frontalpoleL; lateralorbitofrontalL; medialorbitofrontalL; parsorbitalisL |
| RPF | Right Prefrontal | frontalpoleR; lateralorbitofrontalR; medialorbitofrontalR; parsorbitalisR |

**Supplementary Table 3. Desikan-Killiany (DK) Cortical Atlas Regions.** Associated two-letter abbreviations for DK cortical atlas regions^64^, and contributing cortical nodes. L/R designation at the end of DK Atlas Node names refer to Left and Right respectively.

| **Feature** | **Comparison** | **Effect Size** | **95% CI** |
| --- | --- | --- | --- |
| Amplitude | Group (Overall) | d = -0.47 | [-0.51, -0.43] |
| Amplitude | Sex | d = 0.38 | [0.33, 0.42] |
| Period | Group (Overall) | d = -0.60 | [-0.64, -0.56] |
| Burst Count | Group (Overall) | RR = 1.11 | [1.04, 1.18] |
| Burst Count | Group (Males) | RR = 1.19 | [1.09, 1.29] |
| Burst Count | Group (Females) | RR = 1.03 | [0.94, 1.13] |

**Supplementary Table 4. Effect sizes for whole-brain alpha burst feature comparisons.** Effect sizes are reported as Cohen's d [95% CI] for amplitude and period, and Rate Ratio [95% CI] for burst count. Negative Cohen's d indicates FXS > TDC. Rate Ratio > 1 indicates TDC > FXS.

| **Region** | **Amplitude (d)** | **Period (d)** | **Burst Count (RR)** |
| --- | --- | --- | --- |
| L Occipital | -0.73 [-0.90, -0.56] | -0.34 [-0.51, -0.18] | 1.10 [1.01, 1.18] |
| R Occipital | -0.71 [-0.88, -0.54] | -0.35 [-0.52, -0.18] | 1.13 [1.04, 1.22] |
| L Temporal | -0.74 [-0.85, -0.63] | -0.38 [-0.49, -0.28] | 1.12 [1.04, 1.21] |
| R Temporal | -0.79 [-0.89, -0.68] | -0.39 [-0.50, -0.29] | 1.13 [1.05, 1.22] |
| L Parietal | -0.56 [-0.73, -0.40] | -0.76 [-0.93, -0.59] | 1.16 [1.08, 1.26] |
| R Parietal | -0.66 [-0.83, -0.49] | -0.70 [-0.87, -0.53] | 1.19 [1.10, 1.28] |
| L Central | -0.45 [-0.65, -0.26] | -0.91 [-1.11, -0.71] | 1.11 [1.03, 1.20] |
| R Central | -0.53 [-0.73, -0.34] | -0.74 [-0.94, -0.54] | 1.14 [1.06, 1.23] |
| L Cingulate | -0.79 [-0.96, -0.61] | -0.80 [-0.97, -0.63] | 1.10 [1.02, 1.19] |
| R Cingulate | -0.81 [-0.98, -0.63] | -0.78 [-0.96, -0.61] | 1.12 [1.03, 1.20] |
| L Frontal | -0.47 [-0.62, -0.32] | -0.69 [-0.84, -0.54] | 1.02 [0.95, 1.10] |
| R Frontal | -0.40 [-0.54, -0.25] | -0.52 [-0.67, -0.37] | 1.03 [0.95, 1.11] |
| L Prefrontal | -0.60 [-0.77, -0.43] | -0.50 [-0.66, -0.33] | 1.07 [0.99, 1.16] |
| R Prefrontal | -0.46 [-0.63, -0.29] | -0.52 [-0.68, -0.35] | 1.08 [1.00, 1.16] |

**Supplementary Table 5. Regional effect sizes for group comparisons.** Effect sizes are reported as Cohen's d [95% CI] for amplitude and period, and Rate Ratio [95% CI] for burst count. Negative Cohen's d indicates FXS > TDC. Rate Ratio > 1 indicates TDC > FXS.

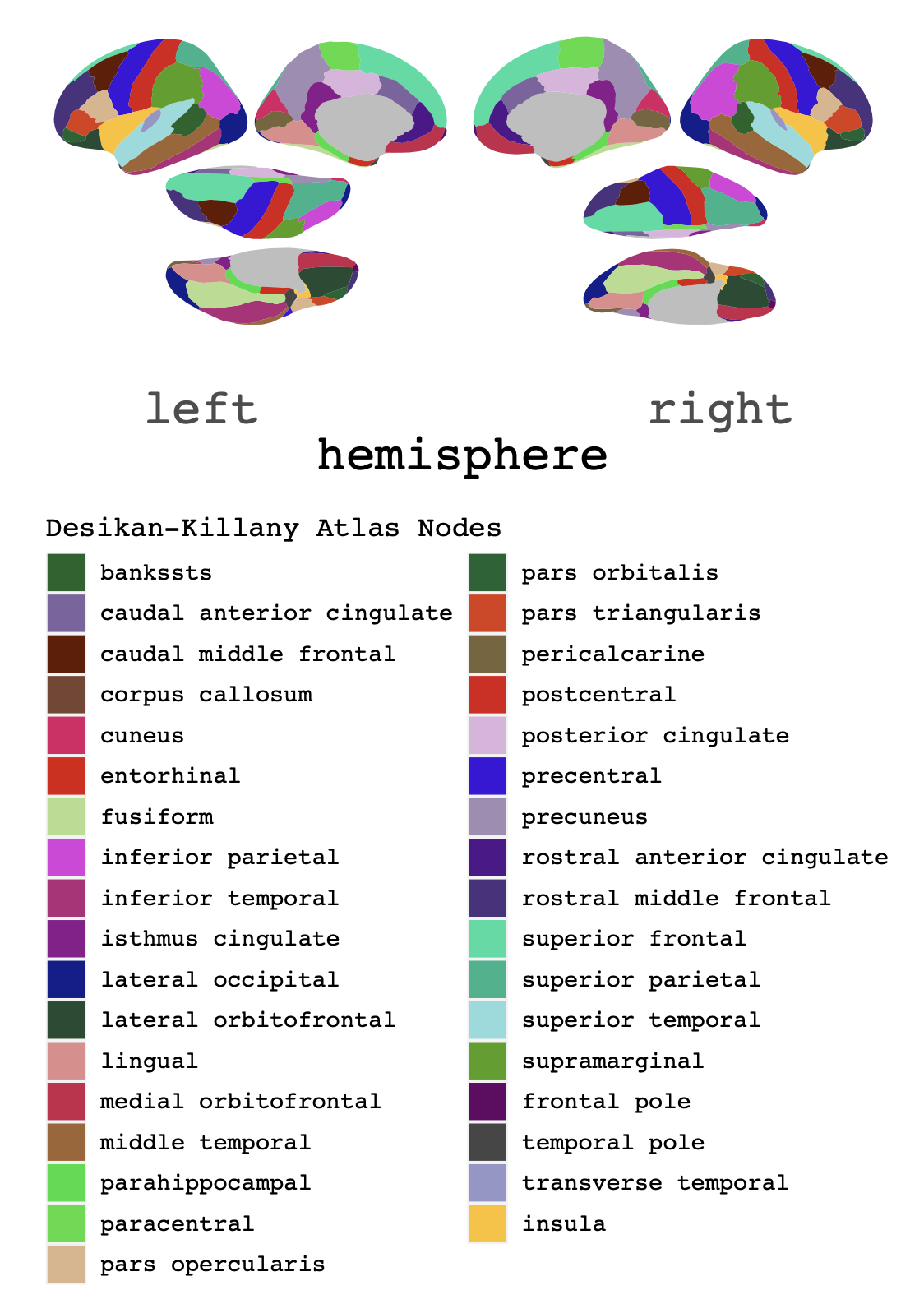
**Supplementary Fig. 1. Source localization parcellation key by node for Desikan-Killiany Cortical Atlas^64^.**

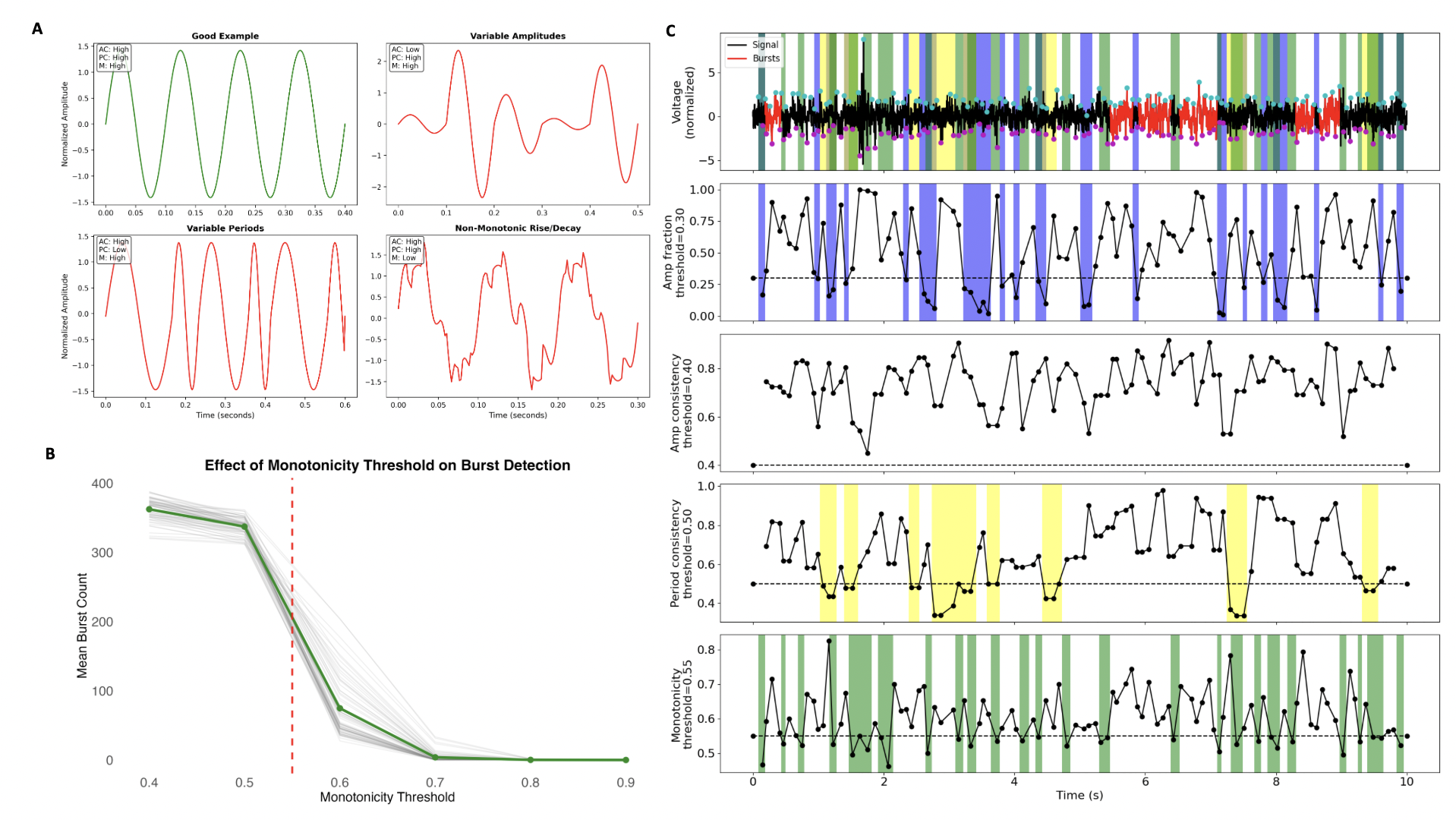

**Supplementary Fig. 2. Monotonicity optimization in alpha burst detection parameter determination.** **(a)** Simulated oscillatory signals demonstrate the effects of burst detection parameters on cycle classification. Top left panel show ideal signals that meet burst criteria with high amplitude consistency (AC), high period consistency (PC), and high monotonicity (M). Top right panel demonstrates low AC, bottom left shows low PC, and bottom right shows low M. **(b)** Systematic evaluation of monotonicity threshold effects on burst detection sensitivity. Mean burst count averaged across all subjects for all 68 nodes decreases sharply above monotonicity = 0.55 (vertical dashed line), with minimal detection occurring at higher thresholds. This inflection point represents a balance between detection sensitivity and shape constraint specificity for alpha oscillations. **(c)** Final optimized burst detection parameters applied to example alpha signal. Top panel displays raw alpha-filtered signal (black) with identified bursts highlighted in red, overlaid with all detection criteria (colored bars). Lower panels show individual parameter traces over time, with colored backgrounds indicating when each threshold criterion is violated: amplitude fraction (blue), period consistency (yellow), and monotonicity (green). Amplitude consistency was above threshold in this example so no associated color. The integrated approach ensures detection of genuine oscillatory activity while maintaining stringent waveform shape requirements.
